# Leishmanial GP63 acts as a protease for the small pore forming toxin aerolysin

**DOI:** 10.64898/2026.07.29.741567

**Authors:** Chaitanya S. Haram, Sebastian Salinas, Salma Waheed Sheikh, Kai Zhang, Peter A. Keyel

## Abstract

The eukaryotic pathogen *Leishmania major* causes disfiguring cutaneous lesions, whose resolution can be complicated by secondary bacterial infections. Bacteria, including *Aeromonas* spp., also interact with *L. major* promastigotes in the sandfly midgut. The mechanisms by which *L. major* competes with bacteria and resists their toxins are poorly defined. Prior work proposed that *L. major* resists the *Aeromonas*-produced pore-forming toxin aerolysin using an altered GPI-anchor. However, we found that *L. major* is sensitive to aerolysin. Here, we determined the mechanism by which *L. major* promastigotes are sensitive to aerolysin, using flow cytometry and biochemical approaches to analyze promastigotes genetically deficient in enzymes that produce key membrane components. The virulence factor lipophosphoglycan protected *L. major* from aerolysin cytotoxicity. The metalloproteinase GP63 exerted the necessary furin-like protease activity to activate aerolysin. Leishmanial GPI-anchored proteins were necessary for aerolysin heptamerization and killing of *L. major* promastigotes. Finally, mutation of the GPI-anchor binding domain of aerolysin crippled its cytotoxicity, consistent with its reliance on the GPI-anchor binding site to engage GPI anchors on the surface of *L. major* promastigotes. Taken together, we propose the *L. major* virulence factor lipophosphoglycan defends against pore-forming toxins made by bacterial competitors, while the GP63 metalloproteinase activates pro-aerolysin like furin. Overall, this study highlights approaches microbes use to compete with each other.

**Graphical Abstract:** **Aerolysin cytotoxicity depends on gp63 and LPG in *Leishmania major* promastigotes.** (A) Wild type *Leishmania major* promastigotes are sensitive to aerolysin, which forms lethal heptameric pore complexes in the plasma membrane (B) *L. major lpg1^—^* promastigotes are highly sensitive to aerolysin challenge because they lack LPG. (C) *L. major gp63^—^* promastigotes have wild type sensitivity to aerolysin challenge but resist pro-aerolysin. (D) *L. major gpi8^—^* knockout promastigotes are resistant to aerolysin and show no heptameric pore complexes in the plasma membrane. Created in BioRender.

## Introduction

Approximately two million new infections annually are attributed to the neglected tropical disease leishmaniasis, caused by protozoan parasites in the genus *Leishmania* (1). Two manifestations of leishmaniasis are disfiguring cutaneous, and mucocutaneous lesions, both of which can become infected with secondary bacterial infections (1). Bacteria also compete with *Leishmania* spp. in the midgut of the sandfly vector. A wide range of bacterial species may compete with *Leishmania* spp., including *Streptococcus*, *Bacillus*, *Flavimonas*, *Pseudomonas*, and *Aeromonas* (2–4). While some of these bacteria may influence lesion pathology or host immune responses, the nature of their interactions with *Leishmania*, especially *Aeromonas* spp., remains poorly understood.

Many bacteria produce pore-forming toxins as virulence factors. Our prior work showed that one subset of bacterial toxins, the cholesterol-dependent cytolysins (CDCs), produced primarily by Gram positive bacteria like *Streptococci*, could only kill *L. major* in the absence of sphingolipids (5). It is not clear if this sphingolipid-based protection is specific to the two CDCs used, streptolysin O (SLO) and perfringolysin O (PFO), all CDCs, or most pore-forming toxins. A second family of bacterial toxins are the aerolysins. The archetypal member of this family, aerolysin, is produced by the Gram negative facultative anaerobes *Aeromonas hydrophila* and *A. salmonicida* (6). Aerolysin is secreted as the inactive precursor pro-aerolysin. After cleavage by furin-like enzymes in the activation loop, aerolysin heptamerizes and inserts into the membrane (7). Aerolysin binds to both the GPI-anchors and N-linked glycans in proteins via two distinct protein domains (8). Glycan binding is disrupted in aerolysin^W45A^, while GPI-anchored binding is eliminated in aerolysin^H332N^ (8). Therefore, aerolysin is likely more influenced by GPI-anchored proteins in *L. major* than by sphingolipids.

*L. major* produces many GPI-anchored molecules. Two prominent GPI-anchored virulence factors are lipophosphoglycan (LPG) and GP63, which play key roles in immune evasion and parasite survival (1). LPG enables host immune evasion via inhibition of maturation of phagosomal lysosome, host adhesion and shields the parasite from stressors (9). GP63 is a well characterized zinc metalloproteinase that cleaves mammalian antimicrobial peptides (AMPs) and mammalian proteins including protein tyrosine phosphatase non-receptor type 12 (PTPN12), mammalian target of rapamycin (mTOR), p65RelA, c-Jun, vesicle-associated membrane protein (VAMP) 3, VAMP8, Nod-like receptor family, pyrin domain-containing 3 (NLRP3), Src homology region 2 (SH2) domain-containing phosphatase 1 (SHP-1), Synaptotagmin XI, and Syntaxin 5 (10). There are also many other GPI-anchored glycoconjugates including GP46, phosphoglycans (PGs), proteophosphoglycans (PPGs), and glycoinositolphospholipids (GIPLs). As in mammalian cells, GPI-anchored proteins can be eliminated in *L. mexicana* by disruption of the transamidase GPI8 (11). However, GPI8 is not needed for LPG synthesis (11). Interestingly, prior work (12) suggested that aerolysin is unable to bind leishmanial GPI anchors. When GP63 was expressed in Chinese Hamster Ovary (CHO) cells, it was recognized by aerolysin, whereas leishmanial lysates containing GP63 were not recognized. (12). However, the ability of aerolysin to kill either *L. major* or CHO cells expressing GP63 was not determined. Thus, the molecular mechanisms by which *L. major* promastigotes resist aerolysin-mediated cytotoxicity remain largely uncharacterized.

In this study, we investigated aerolysin-mediated cytotoxicity in *Leishmania major* promastigotes using flow cytometry and western blots. In contrast to prior findings, we observed killing of *L. major* by aerolysin. We found that LPG was the primary means of resistance to aerolysin, and that aerolysin needs to engage GPI-anchored proteins for binding and cytotoxicity. Surprisingly, we found that leishmanial GP63 can activate pro-aerolysin. Together, our findings provide molecular insight into interdomain competition between *L. major* and bacteria.

## Results

### LPG protects *Leishmania* from aerolysin cytotoxicity

Since loss of SPT2 (required for sphingolipid synthesis) sensitizes *L. major* promastigotes to CDCs (5), we tested if it also sensitized promastigotes to aerolysin. We challenged wild type*, spt2^—^* and *spt2^—^*/+SPT2 promastigotes with either pro-aerolysin or aerolysin. In mammalian cells, pro-aerolysin is activated by furin proteases on the cell surface (7). Aerolysin can also be activated by trypsin prior to cell challenge (13). We used our well-characterized membrane integrity assay to measure killing (5, 14–17). We measured *L. major* killing across a range of toxin concentrations and calculated the amount of toxin needed to kill 50% of the cells, termed LC_50_. Both pro-aerolysin and aerolysin killed promastigotes to a similar extent, indicating that the membrane of *L. major* contains aerolysin-activating activity (Supplementary fig S1A, B). The *spt2^—^* promastigotes were killed to the same extent as wild type and *spt2^—^*/+SPT2 promastigotes (Supplementary Fig S1A, B). Thus, SPT2 failed to protect *L. major* promastigotes from aerolysin, which can be activated by leishmanial surface enzymes.

These findings contrasted with prior results (12), so we explored other potential binding partners for aerolysin in *L. major*. We tested the impact of altering leishmanial ergosterol on aerolysin cytotoxicity using genetic mutants lacking either *sterol methyltransferase* (*smt*), or *C14 demethylase* (*c14dm*). The *smt^—^* promastigotes accumulate cholestane-based sterols like 14-methylated ergosterol (18), while *c14dm^—^* promastigotes accumulate 14-methylated and ergostane-like sterols molecules (19). We challenged *smt^—^* and *c14dm^—^ L. major* promastigotes with pro-aerolysin and aerolysin, and compared their viability to wild type, *smt^—^*/+SMT, and *c14dm^—^*/+C14DM promastigotes. The *smt^—^* promastigotes had wild type sensitivity to pro-aerolysin and aerolysin challenge (Supplementary Fig S1C, D). In contrast, *c14dm^—^* promastigotes were more sensitive to aerolysin than controls (Supplementary Figure S1E, F). These results are consistent with the generally increased sensitivity of *c14dm^—^* promastigotes to detergents and other membrane stress (5, 20). We conclude that ergosterol is not a major contributor to aerolysin sensitivity.

In addition to differences in sterol composition, *c14dm^—^*promastigotes have higher gp63 expression, and lower LPG expression compared to wild type (19). Since aerolysin binds to GPI-anchored species in mammals (21), it is possible that GPI-anchors in *L. major* are masked by some other species, like LPG. We tested the contribution of LPG using the previously characterized LPG null mutant, *lpg1^—^* (22). The *lpg1^—^*promastigotes lack the key galactofuranosyltransferase needed to synthesize the LPG glycan core (22). The *lpg1^—^* promastigotes were two orders of magnitude more sensitive to both pro-aerolysin and aerolysin challenge than wild type or *lpg1^—^*/+LPG1 promastigotes (Fig 1A, Supplementary Fig S2A). To determine the basis of this sensitivity, we measured aerolysin oligomerization into heptamers by western blot. Aerolysin heptamers formed after pro-aerolysin or aerolysin challenge in all three genotypes tested (Fig 1B), though there was an increase in heptamers in the *lpg1^—^* promastigotes (Fig 1B). This suggests that loss of LPG1 promotes a more permissive environment for aerolysin binding and pore formation.

**Figure 1.**
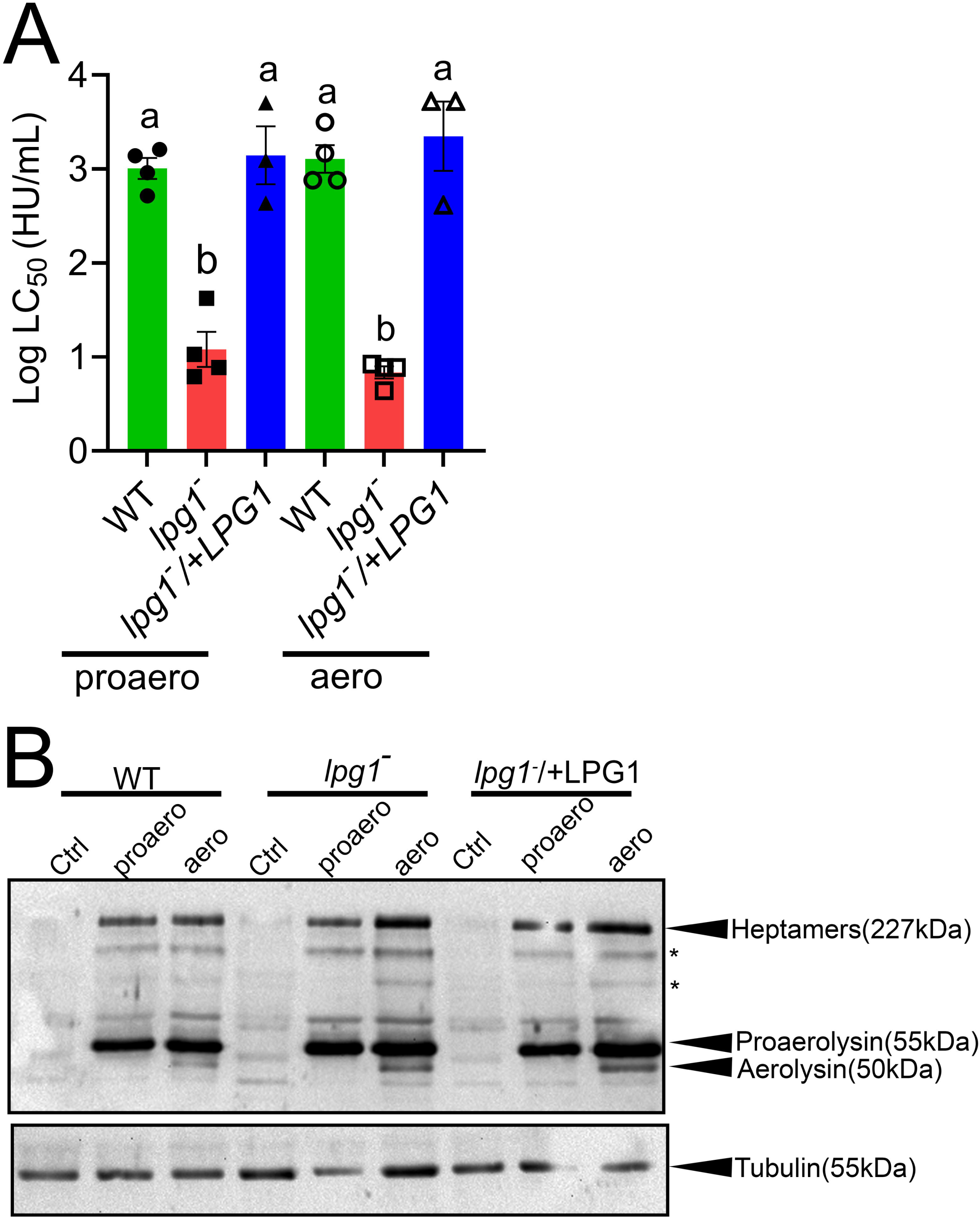
LPG restricts aerolysin binding and cytotoxicity. Wild type (WT), *lpg1^—^* and *lpg^—^* /+LPG *L. major* promastigotes were challenged at 37°C for 1 h with either (A) 31-4000 HU/mL pro-aerolysin (proaero) or aerolysin (aero) or (B) with 1 μg (equivalent to 2000 HU/mL) pro-aerolysin or aerolysin. (A) Cytotoxicity was measured by flow cytometry. The LC_50_ was calculated as described in the methods. (B) Promastigotes were collected in SDS sample buffer, and analyzed by western blot for the indicated proteins. Graphs display at least 3 independent experiments, along with the mean ± s.d. Western blots represent 2 independent experiments. Asterisks denote non-specific bands. Groups sharing the same letter were not statistically different by one way ANOVA with Tukey post hoc testing.

### Leishmanial GP63 activates aerolysin

We next investigated the GPI-anchored proteins to determine the aerolysin binding partner. Since prior work proposed that the leishmanial GPI-anchor of GP63 is in the wrong conformation to sustain aerolysin binding (12), we first tested the impact of deleting *gp63*. We compared the previously characterized *gp63^—^* and *gp63^—^*/+GP63 promastigotes (23) to their wild type strain (Seidman) and the wild type strain used for our other knockouts (LV39). To our surprise, g*p63^—^* promastigotes were resistant to pro-aerolysin, but had wild type sensitivity to trypsin-activated aerolysin (Fig 2A, Supplementary Fig S2B). Consistent with this finding, the *gp63^—^*/+GP63 promastigotes were more sensitive to pro-aerolysin than wild type, but had wild type sensitivity to active aerolysin (Fig 2A, Supplementary Fig S2B). We next tested if this was related to binding or oligomerization. Following active aerolysin challenge, heptamers formed in all genotypes tested, independent of GP63 expression (Fig 2B). However, pro-aerolysin only formed heptamers when GP63 was present (Fig 2B). These data suggest that GP63 is dispensable for aerolysin binding, but is involved in activating pro-aerolysin.

**Figure 2.**
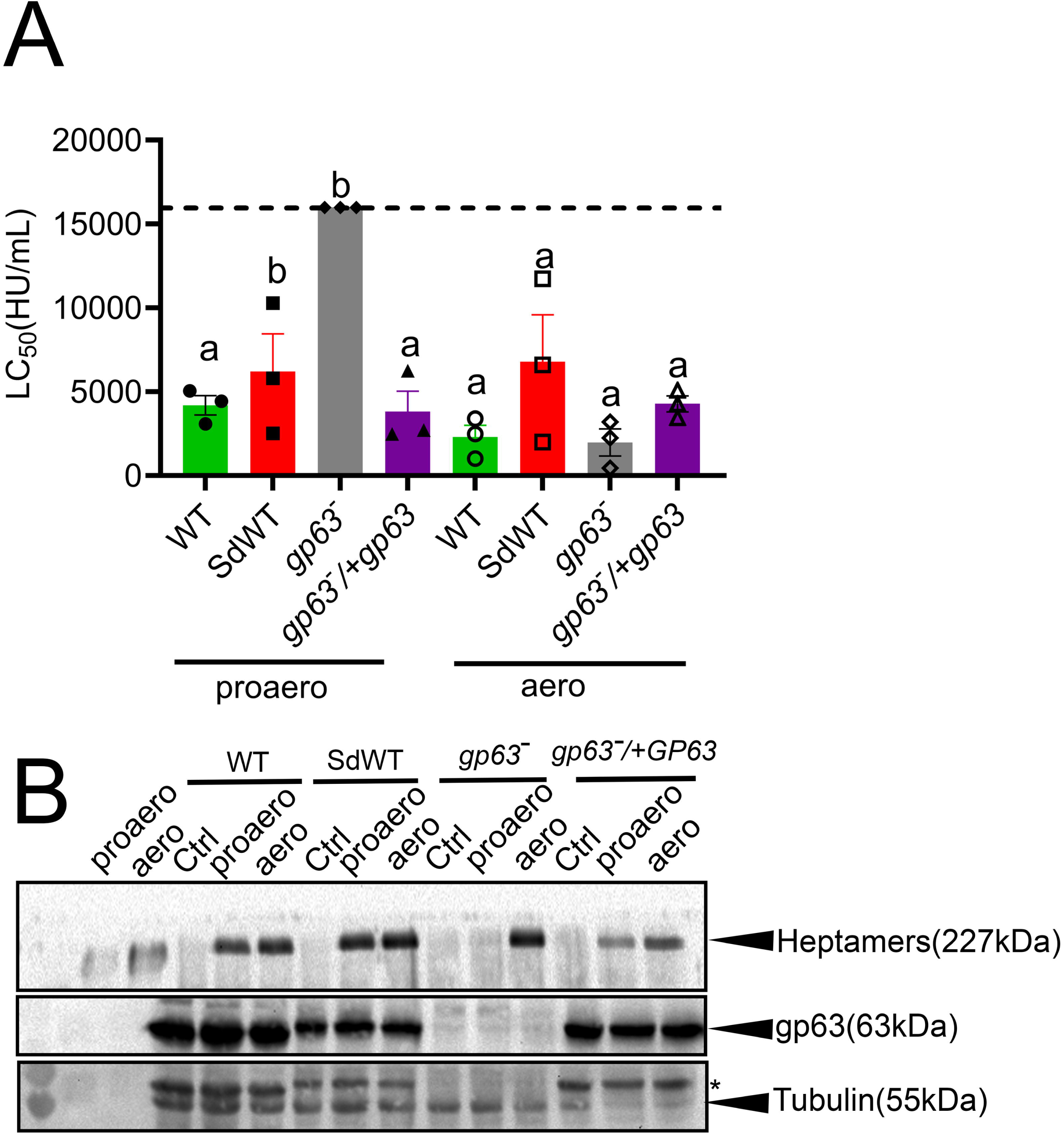
GP63 activates aerolysin. (A, B) Wild type (LV39WT, SDA2WT), *gp63^—^* and *gp63^—^* /+GP63 *L. major* promastigotes were challenged at 37°C for 1 h with either pro-aerolysin (proaero) or aerolysin at a maximum dose of 6000HU/mL or (B) 1 μg pro-aerolysin or aerolysin. (A) Cytotoxicity was measured by flow cytometry. The LC_50_ was calculated as described in the methods. (B) Promastigotes were collected in SDS sample buffer, and analyzed by western blot for the indicated proteins. (C-F) Lysates from the indicated genotypes of *L. major* were incubated with pro-aerolysin for 60 min at 37° C. Aerolysin activation and gp63 levels were measured by western blot, using purified pro-aerolysin or trypsin-activated aerolysin as controls. The ratio of pro-aerolysin to active aerolysin was determined by densitometry. Graphs display at least 3 independent experiments, along with the mean ± s.d. Western blots represent 2 independent experiments. Asterisks denote incomplete GP63 stripping. Groups sharing the same letter were not statistically different by one way ANOVA with Tukey post hoc testing.

Since GP63 is a metalloproteinase, we next tested the ability of GP63 to contribute the furin-like activity that activates aerolysin using an *in vitro* assay. We incubated pro-aerolysin with membrane lysates from *L. major* promastigotes expressing or not expressing GP63, and measured the conversion of pro-aerolysin to aerolysin. In contrast to controls and *lpg1^—^* promastigotes, lysates from *gp63^—^* promastigotes were unable convert pro-aerolysin to aerolysin (Fig 3). Based on these data, we conclude that GP63 activates aerolysin.

**Figure 3.**
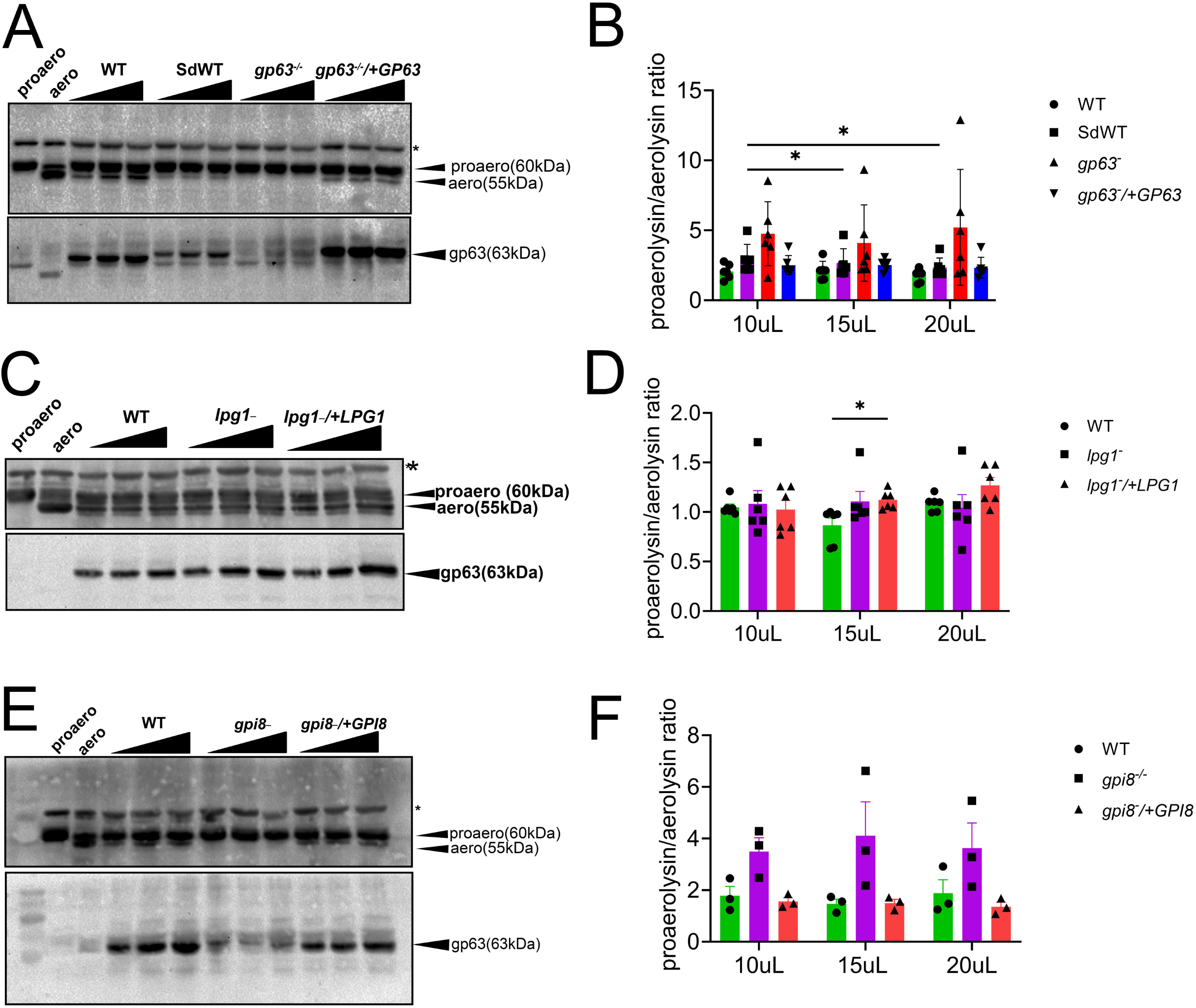
Leishmanial gp63 acts as a novel metalloprotease for activation of aerolysin. (A) LV39WT, Seidmann (SDA2WT), *gp63^-^*and *gp63^-^*/+GP63 or (C) LV39WT, *lpg1^-^* and *lpg^-^*/+LPG (E) LV39WT, gpi8^—^ & gpi8^—^/+GPI8 *L. major* lysates were incubated with 5 μg pro-aerolysin at 37°C for 1hr. Samples were run on a 10% SDS PAGE gel, transferred on nitrocellulose membrane and blotted with antisera raised against aerolysin, gp63, alpha tubulin and Bip. Asterisks denote non-specific bands. (B,D,F) Pro-aerolysin/aerolysin ratio for each volume of lysate calculated from intensities of pro-aerolysin and aerolysin band normalized to its background from at least 3 independent blots for each set of genotypes. * p < 0.05 by one way ANOVA with Tukey post hoc testing.

### Leishmanial GPI-anchored proteins are needed for aerolysin heptamerization and killing

While GP63 is the major GPI-anchored protein in *L. major*, it is not the only one. To test if other GPI-anchored proteins sustained aerolysin binding and pore-formation in the absence of GP63, we eliminated the key transamidase that links proteins to GPI-anchors, *gpi8* (17). Promastigotes lacking *gpi8* were viable, grew as expected, had elevated levels of LPG, but lacked GPI-anchored proteins like GP63 (17, 24). We challenged *gpi8^—^* and *gpi8^—^*/+GPI8 promastigotes with pro-aerolysin or aerolysin. The *gpi8^—^* promastigotes resisted aerolysin independent of its activation status (Fig 4A, Supplementary Fig S2C). We next tested heptamerization, and found that while heptamer formation was sustained in control cells, *gpi8^—^* promastigotes failed to sustain any heptamerization (Fig 4B). We further tested the ability of *gpi8^—^* promastigote lysates to cleave aerolysin. Consistent with the lack of GP63, *gpi8^—^*lysates failed to activate aerolysin (Fig 3E-F). Based on these data, we conclude that GPI-anchored proteins are necessary for aerolysin heptamerization and killing in *L. major* promastigotes.

**Figure 4.**
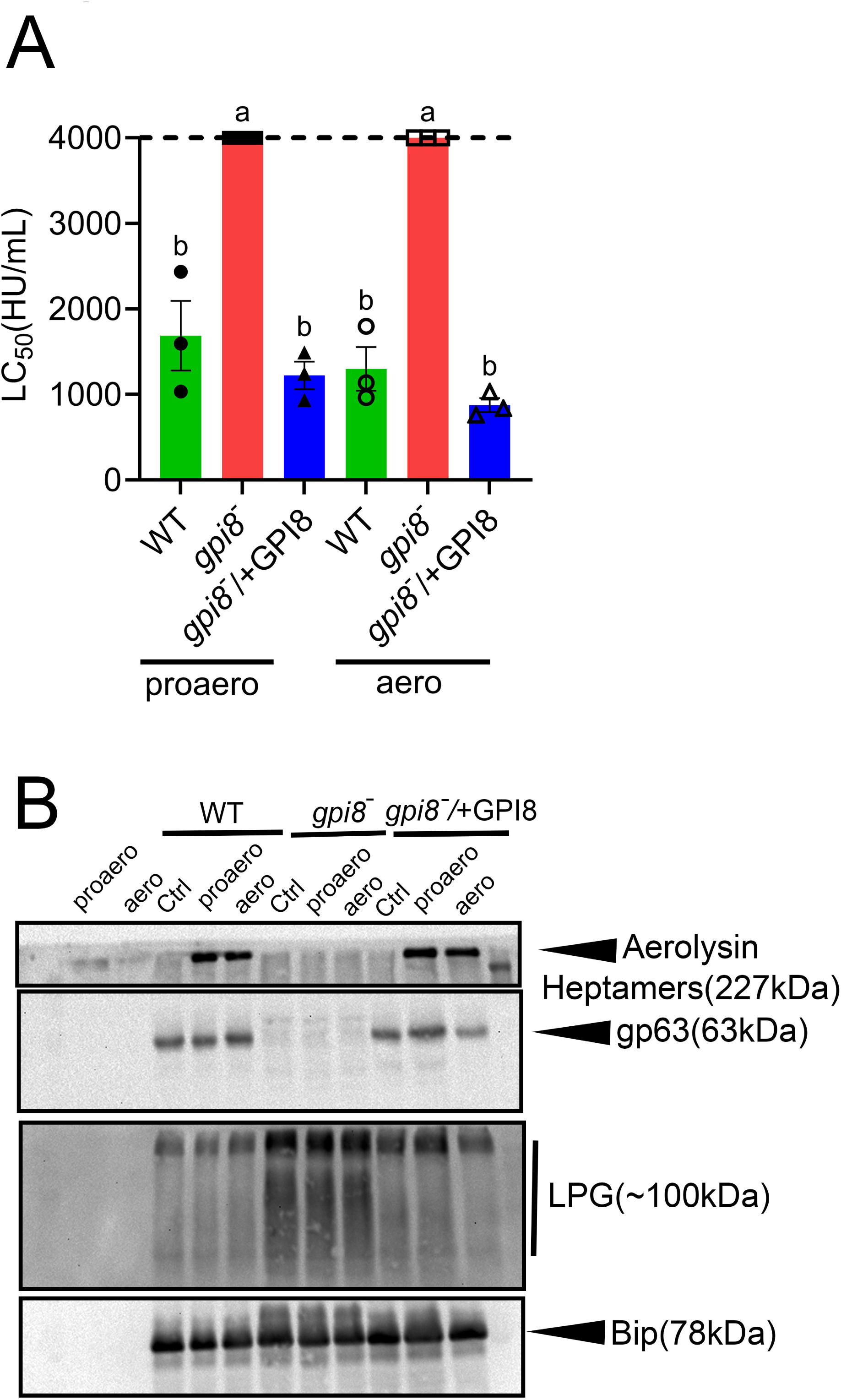
Aerolysin requires the leishmanial GPI-anchor to bind and kill *L. major* promastigotes. (A, B) LV39WT, *gpi8^—^*, and *gpi8^—^*/+GPI8 *L. major* promastigotes were challenged at 37°C for 1 h with either (A) 4000 HU/mL maximum dose of pro-aerolysin or aerolysin or (B) 1 μg of pro-aerolysin and aerolysin at 37°C for 60 min. (B) Promastigotes were subsequently collected in 1X SDS-sample buffer, run on SDS-PAGE, transferred onto nitrocellulose membrane and probed with antisera against aerolysin, LPG and tubulin. (C-D) Lysates from the indicated genotypes of *L. major* were incubated with pro-aerolysin and analyzed as described in Fig 2. Graphs display at least 3 independent experiments, along with the mean ± s.d. Western blots represent 2 independent experiments. Groups sharing the same letter were not statistically different by one way ANOVA with Tukey post hoc testing.

### The GPI-anchor binding domain of aerolysin drives cytotoxicity in *L. major*

To corroborate the need for GPI-anchored proteins in aerolysin killing of *L. major*, we tested the two key binding domains on aerolysin. Aerolysin domain 1 binds to N-linked glycans, while domain 2 engages the GPI anchor (8). These domains can be disrupted by introducing W45A and H332N mutations, respectively, into aerolysin (8). We challenged wild type, *gp63^—^*, *gpi8^—^*, or *lpg1^—^* promastigotes with aerolysin, aerolysin^W45A^, or aerolysin^H332N^. Disruption of either domain increased the LC_50_ for aerolysin in all genotypes tested (Fig 5A, Supplementary Fig S3A-C), indicating both domains contribute to aerolysin cytotoxicity in *L. major*. Elimination of the GPI-anchor binding site (aerolysin^H332N^) crippled cytotoxicity, indicating aerolysin uses this domain during aerolysin cytotoxicity (Fig 5A, Supplementary Fig S3A-C). Since overexpression of GP63 increased killing of *L. major* promastigotes in the absence of the GPI-anchor binding site, aerolysin might engage GP63 via N-linked glycans (Fig 5A, Supplementary Fig S3A-C). As expected, *gpi8^—^* promastigotes remained resistant to all aerolysin variants (Fig 5B-D, Supplementary Fig S3D). In the sensitive *lpg1^—^* promastigotes, disruption of either glycan-binding or GPI-anchor binding increased the aerolysin LC_50_ (Fig 5E, Supplementary Fig S3E-G). However, the LC_50_ remained below that from wild type *L. major* challenged with wild type aerolysin (Fig 5E, Supplementary Fig S3E-G). We interpret these data to indicate that either binding site can drive cytotoxicity in a permissive environment. Overall, we conclude that the GPI-anchor binding domain is a major determinant of cytotoxicity in *L. major* promastigotes.

**Figure 5.**
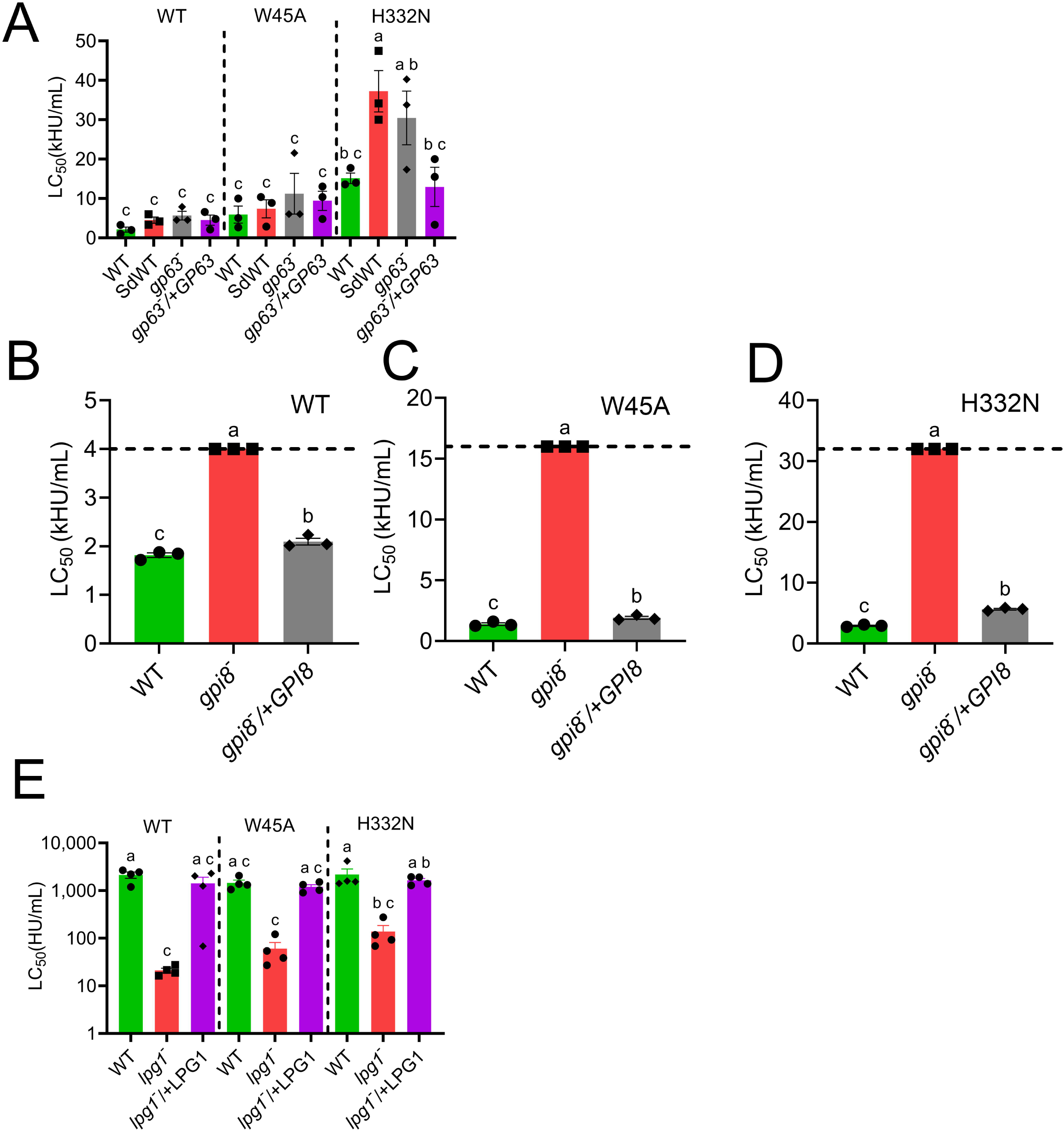
The GPI anchor binding site is needed for aerolysin cytotoxicity. (A) Wild type (LV39WT, SDA2WT), *gp63^—-^* and *gp63^—-^*/+GP63 or (B-D) LV39WT, *gpi8^—^*, and *gpi8^—^*/+GPI8 *L. major* promastigotes were challenged with activated aerolysin (WT), aerolysin^W45A^, or aerolysin^H332N^ at 37°C for 1 h, and analyzed by flow cytometry. LC_50_ was calculated via logistic modeling. Graphs display 3 independent experiments, along with the mean ± s.d. Groups sharing the same letter were not statistically different by one way ANOVA using Tukey post-hoc testing.

## Discussion

Here, we tested the sensitivity of *L. major* to aerolysin, using promastigotes genetically deficient in enzymes needed to produce critical membrane components. We present two key findings: LPG protects *L. major* from aerolysin cytotoxicity instead of altered GPI-anchor conformations, and the metalloproteinase GP63 has the furin-like protease activity needed to activate pro-aerolysin. These findings reveal competitive mechanisms between *Leishmania* spp. and their bacterial competitors.

We found that loss of LPG renders *L. major* ∼100-fold more sensitive to aerolysin, despite no changes to the *L. major* GPI anchor. We interpret these data to indicate that LPG plays a central role in defense against aerolysin, potentially due to its high abundance, larger size (∼20 nm), and shorter lysoalkyl chain positioning LPG close to the surface of the plasma membrane. Since we observed no major changes in aerolysin binding or heptamerization, the mechanism of defense may be interference with pore insertion or limiting pore diffusion over the entire parasite surface. While it is possible that loss of LPG destabilizes the membrane or otherwise makes the promastigotes more fragile, we do not think this is the case. Promastigotes deficient in *lpg1* had no difference in their sensitivity to cholesterol-dependent cytolysins (17). Our present work provides a foundation for future studies examining how pore-forming toxins can be inhibited after oligomerization.

The ability of LPG to provide protection may saturate above wild type levels. Supporting this idea, both the *spt2^—^* and *smt^—^*promastigotes, which have higher LPG levels (18, 25), had similar sensitivity compared to wild type promastigotes. Overexpression of *lpg1* similarly failed to provide additional protection. However, the Seidman strain was more resistant to aerolysin at baseline levels. We attribute differences in the sensitivity of the LV39 and Seidman strains to differences in their respective LPG. Our work provides a foundation for future work into determining the structural features of LPG that prevent aerolysin cytotoxicity.

We found that leishmanial GPI-anchors support binding of aerolysin. This contrasts with a prior report (12) suggesting that the conformation of GPI-anchors in *Leishmania* spp. fail to support binding. We account for these discrepancies in two ways. First, we attribute the membrane environment, especially the presence of LPG, to the differential binding observed by Diep et al. Second, the GP63 tested may have been the catalytically inactive E265D mutant because wild type GP63 could not be stably expressed in CHO cells (26). By contrast, we tested both live promastigotes genetically deficient in key lipid components and aerolysin with either key binding site disrupted. We confirmed that the GPI-anchor binding site on aerolysin is critical to *L. major* cytotoxicity. Like mammalian cells (27), both sites were needed for maximal cytotoxicity. Aerolysin was further unable to bind or kill *gpi8^—^* promastigotes, suggesting that GPI-anchored proteins are needed for binding in *L. major*.

Surprisingly, we found that GP63 can exert the “furin-like activity” needed to activate aerolysin. Furin cleaves after its consensus sequence RXXR (28). Furin activates aerolysin by cleaving after the sequence KVRRAR_432_ in the activation loop of domain 4 (7). In contrast, GP63 cleaves prior to□[K/R][K/R] (29). Since this sequence is also present in the activation loop (VRR), it is possible that GP63 cuts after K427 instead of R432. We expect GP63 to cleave within the activation loop because GP63-activated aerolysin showed similar killing to trypsin-activated aerolysin, and the sizes of the active aerolysins were similar by SDS-PAGE. Our finding adds aerolysin to the list of proteins cleaved by GP63, though this cleavage may be deleterious to the parasite. Thus, we expand the types of protease that can activate aerolysin.

While we found contrasting roles for LPG and GP63 in resisting aerolysin, our study had limitations. The contributions of minor GPI-anchored glycoconjugates such as PGs, PPGs, and GIPLs remain unexplored. The distinct functions of the structural domains of LPG require further analysis using targeted mutants like *lpg2^—^*, *lpg3^—^*, and *lpg5^—^*. While we examined many membrane components, we did not test all of them. We did not identify the direct binding partner for aerolysin using assays such as pull-downs, potentially due to multiple potential binding partners, and complex, or low-abundance components. Similarly, we did not directly quantify cell binding or oligomerization for aerolysin^W45A^ or aerolysin^H332N^ in *L. major* promastigotes, so reduced cytotoxicity could reflect altered post-binding steps. Identifying which species facilitate toxin oligomerization is a new direction opened by this research.

Overall, our work provides insight into competition between *Leishmania* spp and bacteria, and interaction between their respective virulence factors. Aerolysin can bind to leishmanial GPI-anchors, and contains sequences that enable its activation by the metalloproteinase GP63. *Leishmania* spp. defend themselves from this toxin via their surface LPG.

### Experimental Procedures

#### Reagents

All reagents were from Thermofisher Scientific (Waltham, MA, USA), unless otherwise noted. The pET22b plasmid encoding His-tagged aerolysin was a kind gift from Gisou van der Goot (École Polytechnique Fédérale de Lausanne, Canton of Vaud, Switzerland) (30). This wild type aerolysin contained Q254E, R260A, R449A and E450Q mutations and the C-terminal KSASA sequence was replaced with NVSLSVTPAANQLE HHHHHH compared to sequences in GenBank (M16495.1). The W45A, H332N and K244C mutations were introduced into wild type aerolysin by Quikchange mutagenesis. Rabbit anti-aerolysin antisera was previously described (31). Anti-LPG monoclonal antibody WIC79.3 was from Stephen Beverley (Washington University School of Medicine, St Louis, MO, USA) (32). Anti-GP63 monoclonal antibody m235 was from W. Robert McMaster (The University of British Columbia, Vancouver, BC, Canada) (33). Mouse anti-tubulin antisera was from Fisher. Cross-adsorbed HRP-conjugated anti-mouse IgG and anti-rabbit IgG was from Jackson Immunoresearch (West Grove, PA).

#### Recombinant Toxins

Toxins were induced and purified as previously described (34, 35). Toxins were induced with 0.2 mM IPTG for 3 h at room temperature and purified using Nickel-NTA beads. Protein concentration was determined by Bradford assay. Hemolytic activity was determined as previously described (14, 16, 17) using human red blood cells (Zen Bio, Research Triangle Park, NC, USA). One hemolytic unit is defined as the amount of toxin required to lyse 50% of a 2% human red blood cell solution in 30 min at 37 °C in 2 mM CaCl_2_, 10 mM HEPES, pH 7.4, and 0.3% bovine serum albumin (BSA) in PBS.

#### HeLa cell culture

Hela cells (ATCC (Manassas, VA, USA) CCL-2) were maintained at 37°C, 5% CO_2_ in DMEM (Corning, Corning, NY, USA) supplemented with 10% Equafetal bovine serum (Atlas Biologicals, Fort Collins, CO, USA) and 1× L-glutamine. They were negative for mycoplasma by microscopy.

#### Leishmania strains and culture

LV39 clone 5 (Rho/SU/59/P) was used as the wild type strain, and most genetic mutants were made in this background (Table 2). The *L. major* strain NIH S (MHOM/SN/74/Seidman) clone A2 background was used for GP63 knockouts (23). Promastigotes were cultured at 27°C in M199 medium (Gibco) with 0.182% NaHCO_3_, 40 mM HEPES, pH 7.4, 0.1 M adenine, 1 µg/mL biotin, 5 µg/mL hemin & 2 µg/mL biopterin and 10% heat inactivated fetal bovine serum (FBS), pH 7.4. Episomal complemented cells were maintained in complete medium in the presence of 10 µg/mL geneticin (G418) except experimental passages.

**Table 1.** Specific activity of active toxin preps used.

| Toxin | Figure Used | Specific Activity (HU/mg) |
| --- | --- | --- |
| Aerolysin | 1A, S2A | $2 \times 10^6$ |
| | 2A, S2B | $2.52 \times 10^6$ |
| | 3A,S2C | $7.61 \times 10^5$ |
| | S1A-D | $6.25 \times 10^4$ |
| | S1E,F | $8.37 \times 10^5$ |
| | 5A,B,S3A,D | $1.28 \times 10^6$ |
| Aerolysin<sup>W45A</sup> | 5A,C, S3B,D | $8.77 \times 10^4$ |
| Aerolysin<sup>H332N</sup> | 5A,D, S3B,D | $8.77 \times 10^4$ |

**Table 2.** Leishmania strains used in this study.

| Name | Genotype | Strain | Ref |
| --- | --- | --- | --- |
| *spt2*<sup>−</sup> | Δ*spt2*::HYG/Δ*spt2*::PAC | LV39 | (25) |
| *spt2*<sup>-</sup>/+SPT2 | Δ*spt2*::HYG/Δ*spt2*::PAC/+pXG-SPT2 | LV39 | (25) |
| *smt*<sup>-</sup> | Δ*smt*::HYG/Δ*smt*::PAC | LV39 | (18) |
| *smt*<sup>-</sup>/+SMT | Δ*smt*::HYG/Δ*smt*::PAC/+pXG-SMT | LV39 | (18) |
| *c14dm*<sup>-</sup> | Δ*c14dm*::HYG/Δ*c14dm*::PAC | LV39 | (19) |
| *c14dm*<sup>-</sup>/+C14DM | Δ*c14dm*::HYG/Δ*c14dm*::PAC/+pXG-C14DM | LV39 | (19) |
| *lpg*<sup>-</sup> | Δ*LPG1*::HYG/Δ*LPG1*::PAC | LV39 | (22) |
| *lpg*<sup>-</sup>/+LPG | Δ*LPG1*::HYG/Δ*LPG1*::PAC/+pSNBR-LPG1 | LV39 | (22) |
| *gp63*<sup>-</sup> | Δ*GP63*::SAT/Δ*GP63*::HYR | Seidman | (23) |
| *gp63*<sup>-</sup>/+GP63 | Δ*GP63*::SAT/Δ*GP63*::HYR/+ pLEXNeo-gp63 | Seidman | (23) |
| *gpi8*<sup>-</sup> | Δ*GPI8*:: BSD/ Δ*GPI8*::HYR | LV39 | (17) |
| *gpi8*<sup>-</sup>/+GPI8 | Δ*GPI8*:: BSD/ Δ*GPI8*::HYR/+ pXG1a-gpi8 | LV39 | (17) |

Culture density and cell viability were determined by hemocytometer counting and flow cytometry after propidium iodide (PI) staining at a final concentration of 20 µg/mL. In this study, log phase promastigotes refer to replicative parasites at 2.0 – 8.0 x10^6^ cells/mL.

#### Leishmania processing

Cells were cultured in complete medium to log phase or stationary phase, according to experimental requirements. Cells were counted, centrifuged at 3000 RPM (Rotor SX4750) for 8-10 min to pellet cells at room temperature (25°C). Cells were washed with 1X PBS and counted again for accuracy. Cells were centrifuged again at 3000 RPM (Rotor SX4750) for 8-10 min at room temperature, and resuspended in serum free M199. The final concentration of cells was 1.0 x10^6^ cells/mL or 1.0 X10^7^ cells/mL for cytotoxicity or western blot, respectively.

#### Aerolysin activation assay

Mid-log to late-log phase promastigotes were processed as described above to collect 2.0 X10^7^ cells. Cells were resuspended in 250 µL RIPA buffer (Fisher Catalog 89900) with 0.01 mM PMSF and incubated at 4°C for 10 min. The lysate was clarified by centrifugation at 16,000xg at 4°C for 10 min. Aliquots (10-20 µL) of the lysate were incubated with 5 μg pro-aerolysin for 60 min at 37°C. The reaction was quenched by adding 12.5 µL 4X SDS-PAGE sample buffer and adjusted to a total volume of 50 µL using water. Samples were heated at 95°C for 10 min. Positive and negative controls were 5 μg proaerolysin treated with or without 0.0025% trypsin for 10 min at room temperature (25°C).

#### Flow cytometry cytotoxicity assay

Cytotoxicity assays were performed as described (14), using 1 x10^5^ *L. major* promastigotes in serum-free M199 supplemented with 2 mM CaCl_2_ and 20 μg/mL propidium iodide (PI) or 1 x10^5^ HeLa cells in RPMI supplemented with 2 mM CaCl_2_ and 20 μg/mL PI. Toxins were added in a two-fold serial dilution. Cells were incubated for 1 h at 37°C. PI fluorescence in cells was measured using an Attune flow cytometer. Debris was gated out. Cells exhibiting high PI fluorescence (1-2 log shift) (PI high), or background PI fluorescence (PI neg) were quantified. Specific lysis was determined as follows: % Specific Lysis = (% PI High^Experimental^ - % PI High^Control^) / (100 - %PI High^Control^). The sublytic dose was defined as the highest toxin concentration that gave <20% specific lysis. The LC_50_ was determined by logistic modeling as described (14).

#### SDS-PAGE and western blotting

Leishmania lysates (from 1X10^7^ cells) collected in 1X SDS-sample buffer were loaded and run on a 10% resolving gel using a Mini Protean III (Bio-Rad) at 150 V for 60 min. Proteins were transferred onto nitrocellulose membrane at 110 V for 90 min. Blots were blocked with 5% bovine serum albumin (BSA) in Tris-buffered saline with 0.1% Tween 20 (TBST) at 4°C for 2 hours. Blots were incubated with primary antibodies (anti-aerolysin 1:5000, anti-LPG 1:1000, anti-GP63 1:1000, anti-tubulin 1:1000) in 1% BSA in TBST overnight at 4°C. Blots were then washed thrice with TBST for 10 min each. Secondary antibodies (1:10,000) in 1% BSA in TBST were added for 1 h at room temperature. Blots were washed with TBST thrice for 10 min each, developed with enhanced chemiluminescence reagent (1.25 mM luminol (Sigma), 0.01% H_2_O_2_ (Walmart, Fayetteville, AR), 0.2 mM p-coumaric acid (Sigma), 0.1 mM Tris-HCl, pH 8.4), and imaged on an iBlot (Invitrogen).

#### Statistics

Prism (Graphpad, San Diego, CA), or Excel were used for statistical analysis. Data are represented as mean ± SEM as indicated. The LC_50_ for toxins was calculated by logistic modeling as previously described (14). Statistical significance was determined either by one-way ANOVA with Tukey post-testing, one-way ANOVA (Brown-Forsythe method) with Dunnett T3 post-testing, or Kruskal-Wallis, as appropriate. p < 0.05 was considered to be statistically significant. Graphs were generated in GraphPad and organized in Photoshop (Adobe, San Jose, CA, USA).

This article contains supporting information. AI pre-review conducted by qedscience.com.

## Data and materials availability

All data are available in the main text or the supporting information.

## Acknowledgments

The authors would like to thank members of the Keyel lab for critical review of the manuscript. We thank colleagues for the generous gifts of reagents. We thank the College of Arts & Sciences Microscopy for use of facilities.

## Author Contributions

Conceptualization: CSH

PAK Methodology: CSH, SS, SWS

Investigation: CSH, SS, KZ, PAK

Data Curation: CSH, SS, PAK

Formal Analysis: CSH, SS, SWS, PAK

Supervision: KZ, PAK

Resources: SWS, KZ

Project Administration: PAK

Funding Acquisition: KZ, PAK

Writing—original draft: CSH, SS, SWS, KZ, PAK

Writing—review & editing: CSH, SS, SWS, KZ, PAK

## Abbreviations

ANOVA: analysis of variance
BSA: bovine serum albumin
BSD: blasticidin
c14dm: C14 demethylase
CDC: cholesterol-dependent cytolysin
CHO: Chinese Hamster Ovary
DMEM: Dulbecco’s Modified Eagle Medium
GIPL: glycoinositolphospholipids
GPI: glycosylphosphatidylinositol
HRP: horseradish peroxidase
HU: hemolytic unit
HYG: hygromycin
IPTG: Isopropyl β-D-1-thiogalactopyranoside
LC50: lethal concentration 50%
LPG: lipophosphoglycan
mTOR: mammalian Target of Rapamycin
NLRP: Nod-like receptor family, pyrin domain-containing
NTA: nitrilotriacetic acid
PBS: phosphate-buffered saline
PG: phosphoglycans
PI: propidium iodide
PMSF: phenylmethylsulfonyl fluoride
PPG: proteophosphoglycans
PTPN12: protein tyrosine phosphatase non-receptor type 12
RIPA: radioimmunoprecipitation assay
RPMI: Roswell Park Memorial Institute medium
SHP-1: Src homology region 2 domain-containing phosphatase 1
smt: sterol 24-C methyltransferase
spt2: serine palmitoyltransferase 2
TBST: Tris-buffered saline with 0.1%Tween
VAMP: vesicle-associated membrane protein
WT: wild type

## Supplemental Figures

**Supplementary Figure S1. Resistance to aerolysin is distinct from resistance to cholesterol-dependent cytolysins.** (A-F) LV39WT, *spt2^—^*, *spt2^—^*/+SPT2, *smt^-^*, *smt^—^*/+SMT, *c14dm^—^* and *c14dm^—^*/+C14DM *L. major* promastigotes were challenged with 64-4000 HU/mL pro-aerolysin (proaero) or aerolysin (aero) at 37°C for 1 h and analyzed by flow cytometry. The LC_50_ was calculated via logistic modeling of the dose response curves. Graphs display either independent experiments, along with the mean ± s.d or mean ± s.d. Groups sharing the same letter were not statistically different by one way ANOVA using Tukey post-hoc testing.

**Supplementary Figure S2. GPI-anchored molecules regulate aerolysin cytotoxicity.** (A) wild type (LV39WT), *lpg1^—^* and *lpg1^—^*/+LPG1, (B) wild type (LV39WT, Seidman WT), *gp63^—^* and *gp63^—^*/+GP63 or (C) wild type (LV39WT), *gpi8^—^* and *gpi8^—^*/+GPI8 *L. major* promastigotes were challenged with the indicated concentrations of pro-aerolysin or aerolysin at 37°C for 1 h. PI uptake was measured by flow cytometry. Graphs display mean ± SEM of at least 3 independent experiments.

**Supplementary Figure S3. Aerolysin uses its GPI anchor binding domain for cytotoxicity against *Leishmania major*.** (A-C) Wild type (LV39WT, Seidman WT), *gp63^—^* and *gp63^—^*/+GP63 or (D) LV39WT, *gpi8^—^* and *gpi8^—^*/+GPI8 *L. major* promastigotes were challenged with the indicated concentrations of aerolysin (WT), aerolysin^W45A^, or aerolysin^H332N^ at 37°C for 1 h. PI uptake was measured by flow cytometry. Graphs display mean ± SEM of at least 3 independent experiments.

