## Supplementary figures and images for "Leishmanial GP63 acts as a protease for the small pore forming toxin aerolysin"

### Supplemental Fig S1

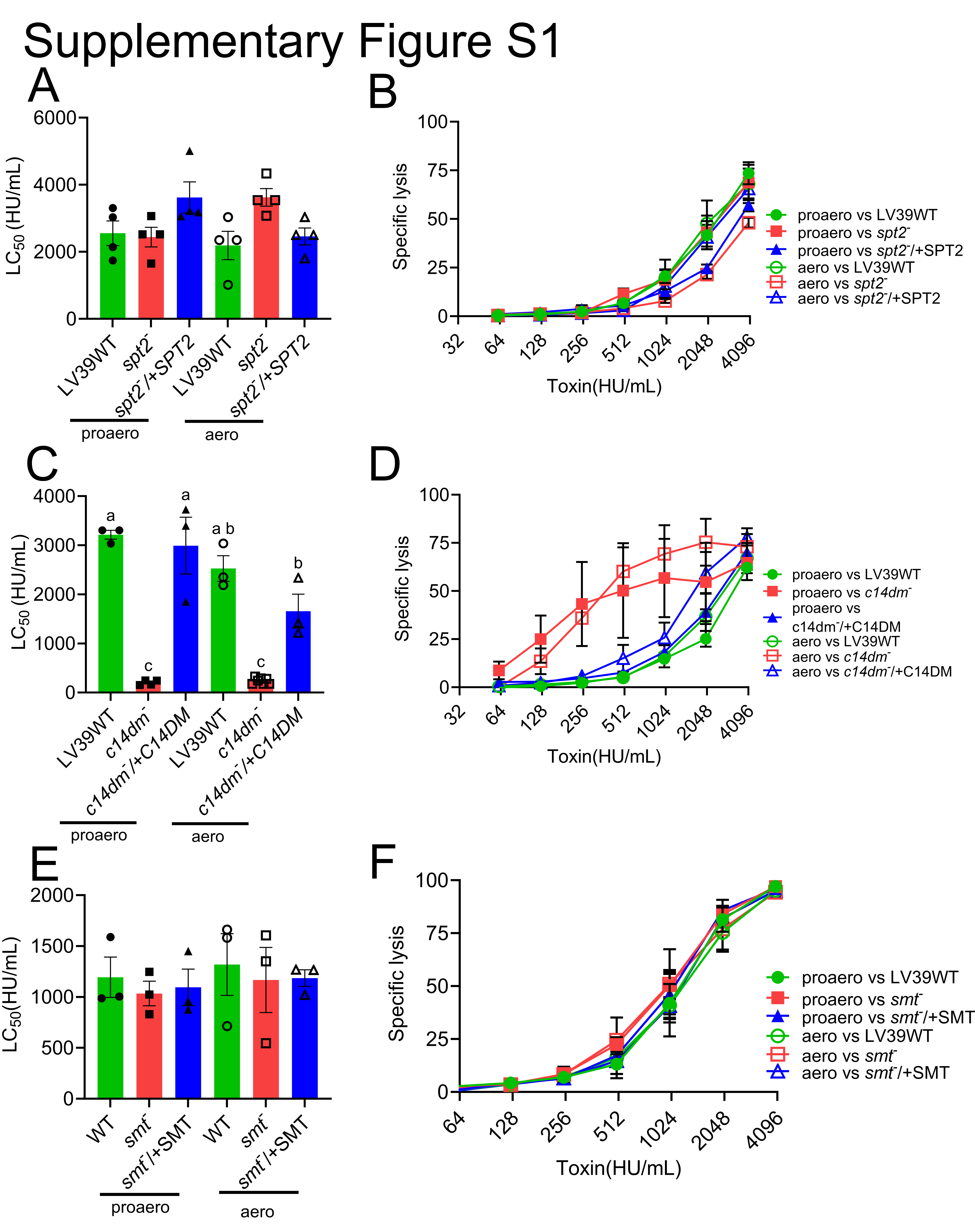

### Supplemental Fig S2

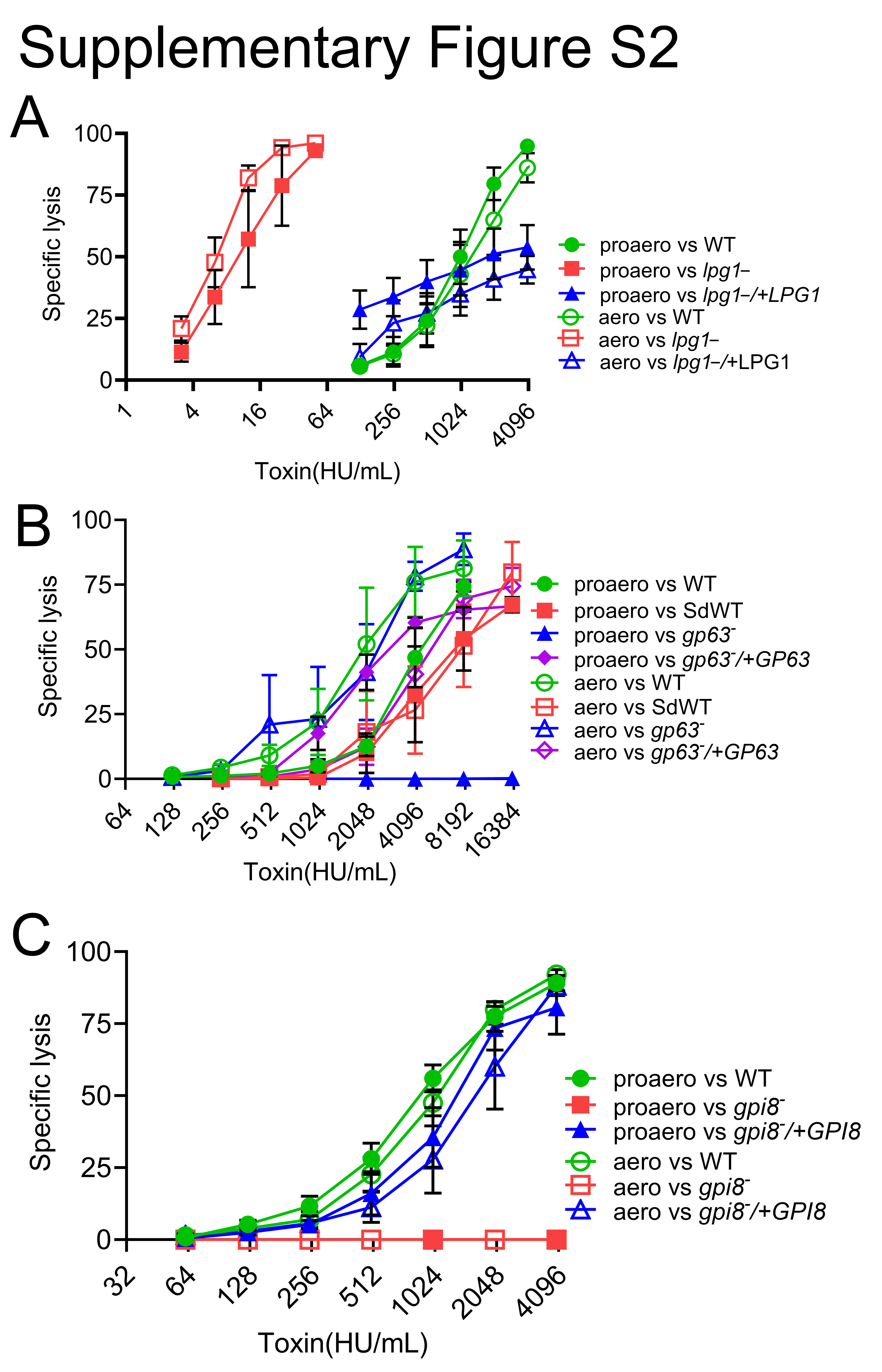

### Supplemental Fig S3

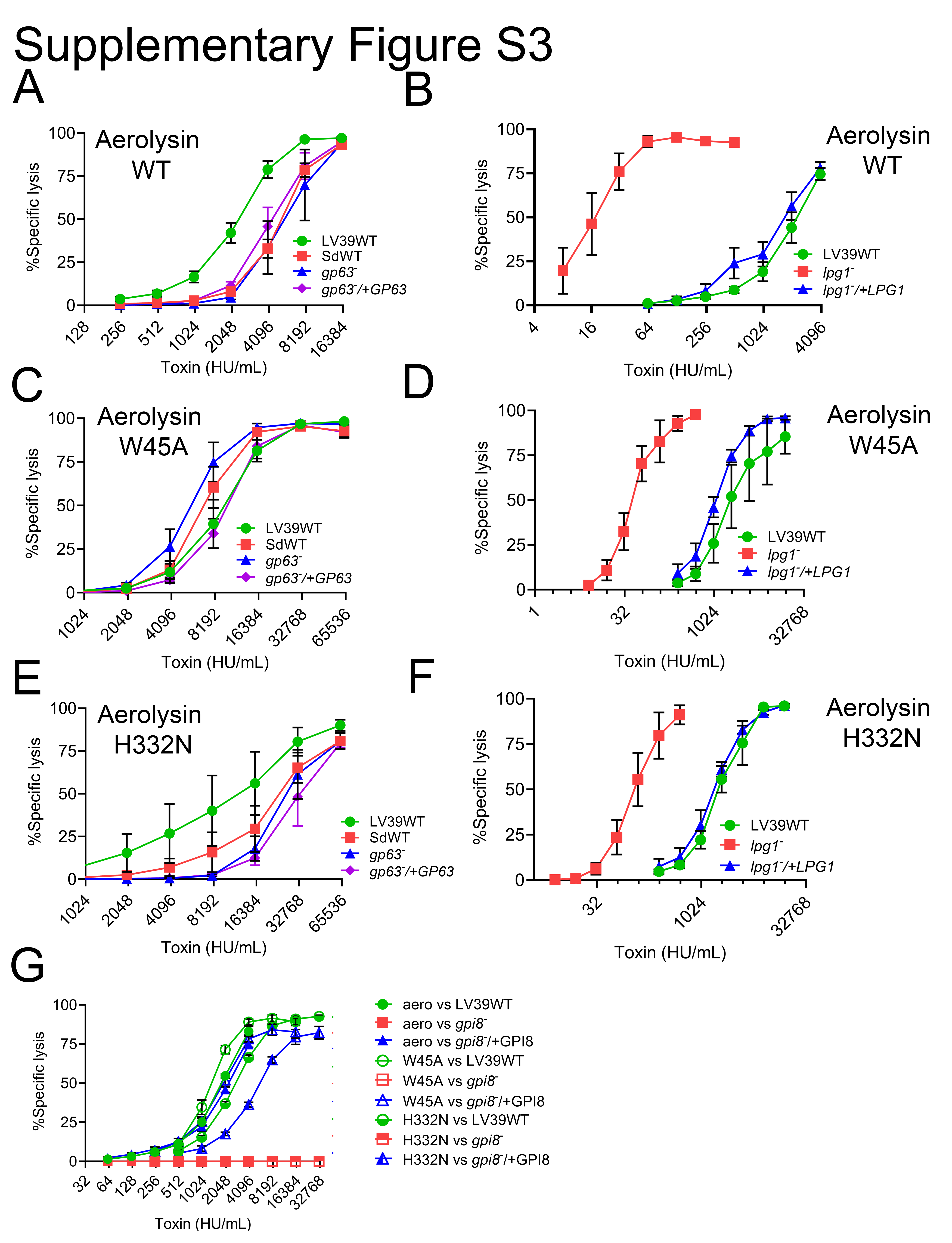
